# Nervous Necrosis Virus Capsid Protein Functions as a SUMO E3 Ligase to Activate MAVS-Dependent NF-κB Signaling

**DOI:** 10.1101/2025.05.29.656765

**Authors:** Wanwan Zhang, Xiaoqi Chen, Bingbing Sun, Lan Yao, Xingchen Xiong, Meisheng Yi, Kuntong Jia

**Affiliations:** School of Marine Sciences, Sun Yat-sen University, Guangzhou 510275, China; Guangdong Provincial Key Laboratory of Marine Resources and Coastal Engineering, Guangzhou 510275, China; Zhuhai Haifa MarineHarvest Industrial Park Co., Ltd. Zhuhai 519000, China

**Author notes:** These authors have contributed equally to this work (Wanwan Zhang, Xiaoqi Chen).

## Abstract

Nervous necrosis virus (NNV) is a lethal aquatic pathogen that activates NF-κB signaling to manipulate host immune responses, yet the underlying molecular mechanisms remain poorly defined. Here, we identify the NNV capsid protein (CP) as a novel viral SUMO E3 ligase that promotes SUMOylation of *Lateolabrax japonicus* mitochondrial antiviral signaling protein (MAVS) to drive NF-κB activation. We show that CP induces p65 nuclear translocation and upregulates proinflammatory cytokine expression in a MAVS-dependent manner. Mechanistically, CP interacts with SUMO2 and the E2 conjugating enzyme *L. japonicus* UBC9 to promote MAVS SUMOylation at lysine 325 (K325), which stabilizes MAVS, facilitates its aggregation, and amplifies downstream NF-κB signaling. Notably, disruption of this modification via the K325R mutation or SUMOylation inhibition abrogates MAVS aggregation and inflammatory responses. Our findings reveal a previously unrecognized strategy by which NNV hijacks the host SUMOylation machinery to fine-tune MAVS function, promoting immune evasion and viral persistence.

**Summary:** Nervous necrosis virus (NNV) is a highly pathogenic virus in aquatic animals that manipulates host immune signaling to promote infection. Here, we identify the NNV capsid protein (CP) as a viral SUMO E3 ligase that modifies the mitochondrial antiviral signaling protein MAVS via SUMOylation at lysine 325. This modification enhances MAVS stability and aggregation, leading to robust activation of the NF-κB signaling pathway and increase expression of proinflammatory cytokines. Disruption of this modification abolishes CP-mediated immune activation. This study unveils a previously unrecognized mechanism by which NNV exploits host SUMOylation machinery to modulate MAVS function, contributing to immune evasion and facilitating viral persistence. These findings provide valuable insights into virus-host interactions and highlight potential therapeutic targets against NNV infections.

## Introduction

Innate immunity represents the first line of defense against viral infections, predominantly triggered by pattern recognition receptors (PRRs) such as RIG-I-like receptors (RLRs) and Toll-like receptors (TLRs) [1]. Once RLRs (e.g., RIG-I and MDA5) detect viral RNA, they transmit signals to the mitochondrial antiviral signaling protein (MAVS), an essential adaptor located on the mitochondrial outer membrane [2]. Activated MAVS subsequently recruits and activates IKK and TBK1, leading to the phosphorylation and nuclear translocation of the transcription factors NF-κB and IRF3, respectively [3]. These transcription factors induce the production of type I interferons (IFN) and pro-inflammatory cytokines, which are vital for the antiviral immune response.

MAVS, as a key activator of NF-κB and IFN pathways, undergoes various regulatory modifications to maintain immune homeostasis, including ubiquitination, phosphorylation, and small ubiquitin-like modifier (SUMO) modification (SUMOylation), all of which significantly influence its signaling capacity and stability [4]. Ubiquitination of MAVS is a well-studied mechanism through which viruses can manipulate host immune responses. For instance, viral proteins of hepatitis C virus (HCV) induce K27-linked ubiquitination of MAVS by the E3 ubiquitin ligase TRIM21, which has a positive regulatory effect on MAVS, thereby promoting the recruitment of TBK1 to MAVS and enhancing downstream signaling [5]. Similarly, phosphorylation of MAVS also regulates its signaling potential, with proteins like VP3 from rotavirus inducing phosphorylation at a newly identified MAVS motif SPLTSS (residues 188– 193), which leads to subsequent K48-linked ubiquitination and degradation [6]. Unlike the transient effects of ubiquitination/phosphorylation, SUMOylation is a dynamic, reversible modification that plays a unique and critical role in the regulation of MAVS activity [7]. This modification has been implicated in enhancing the stability, expression, and function of MAVS during immune responses. For instance, SARS-CoV-2 viral protein Nsp5 advanced the SUMOylation of MAVS to enhance its stability and expression, ultimately activating NF-κB signaling pathway [8]. Similarly, cytosolic poly(dA:dT) DNA promotes SUMO3 conjugation to MAVS, enhancing its aggregation and upregulating IFN-β expression in human keratinocytes [9]. Protein inhibitors of activated STAT3 (PIAS3) and SENP1 (Sentrin/SUMO-specific protease 1) regulate MAVS phase separation in a SUMOylation-dependent manner [10]. These studies highlight the critical role of SUMOylation in shaping MAVS-mediated immune responses, although the precise mechanisms, particularly during viral infections, remain incompletely understood.

Nervous necrosis virus (NNV), classified as *Betanodavirus*, leads to a highly fatal and infectious fish nervous necrosis disease and significant economic losses [11]. The virus genome contains two single-strand RNAs: RNA1 encoding RNA-dependent RNA polymerase (RdRp) and RNA2 encoding the multifunctional capsid protein (CP) [12]. Like other RNA viruses, NNV is recognized by host RLRs (RIG-I/MDA5) that activate MAVS signaling, triggering IRF3/IRF7-mediated type I IFN production and NF-κB-driven inflammatory responses to establish antiviral defense [13, 14]. While studies confirm NNV infection activates NF-κB signaling-with proteasome subunit beta type-8 being identified as a negative regulator of this process [15], the precise mechanism by which NNV subverts the MAVS-NF-κB signaling axis remains unknown. Notably, The NNV CP, as the sole structural component of the virion, plays a crucial role not only in viral assembly and host cell entry but also in modulating the host’s innate immune response [16, 17]. It targets host E3 ubiquitin ligases (RNF34/RNF114) to degrade key RLRs pathway components [18, 19], induces incomplete autophagy via HSP90ab1-AKT-MTOR axis suppression [17], and cooperates with RdRp to disrupt cGAS-mediated IFN signaling [20]. Although these findings establish CP as a master regulator of IFN responses, its potential involvement in NF-κB signaling regulation has never been explored.

In this study, we demonstrate that NNV activates the NF-κB signaling pathway by inducing *Lateolabrax japonicus* MAVS SUMOylation. Specifically, we show that the NNV CP functions as a SUMO E3 ligase, facilitating SUMOylation of MAVS at Lys325. Our findings reveal a previously unrecognized role for the NNV CP in modulating MAVS feedback regulation and NF-κB signaling activation, providing new insights into virus-host interactions and the fine-tuning of MAVS function during viral infection.

## Results

### NNV CP induces NF-κB signaling activation in a MAVS-dependent manner

NNV infection is known to induce an acute inflammatory response in infected fish, with NF-κB signaling playing a central role in this process [13]. To investigate the involvement of NNV CP in NF-κB activation, we assessed the ability of CP to activate this pathway. As shown in Fig 1A, CP overexpression significantly increased NF-κB luciferase activity. Consistent with transcriptional activation, western blot analysis further confirmed that CP overexpression resulted in increased level of phosphorylated p65 (p-p65) in a dose-dependent manner (Fig 1B). Immunofluorescence assays demonstrated that CP promoted nuclear translocation of p65, with p65 localized in the nucleus of CP-expressing cells, in contrast to its cytoplasmic localization in control cells (Fig 1C). These results demonstrate that CP activates NF-κB signaling via p65 nuclear translocation.

**Fig 1.**
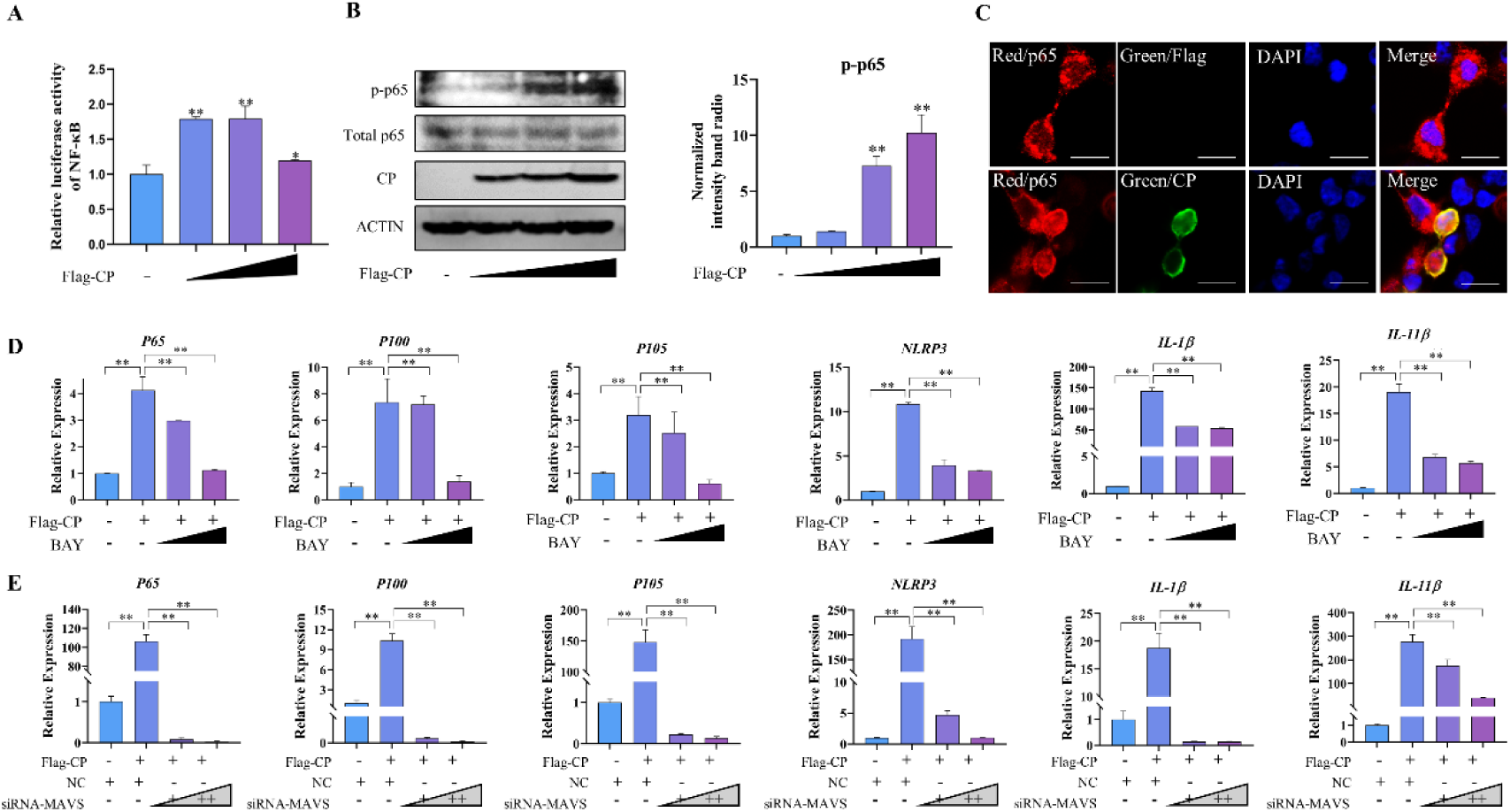
NNV CP activates the NF-κB pathway through a MAVS-dependent manner. (A) Effect of CP on NF-κB activation. HEK 293T cells were co-transfected with pGL3-NF-κB-Luc-pro reporter plasmid (0.5 µg) and varying amounts of *pCMV-Flag-CP* (0, 1, 2, or 3 µg). Luciferase activity was measured at 24 h post-transfection. (B) Dose-dependent effect of CP on p65 protein expression. HEK 293T cells were transfected with increasing doses of *pCMV-Flag-CP* (0, 1, 2, or 3 µg) for 24 h. Protein levels of CP, phosphorylated p65 (p-p65), total p65, and β-actin were detected. (C) Subcellular localization of p65 and CP proteins. The distribution of p65 (red) and CP (green) in HEK 293T cells was analyzed by fluorescence microscopy. Nuclei were stained with DAPI (blue). Scale bar = 10 µm. (D) Effect of CP and Bay 11-7082 (BAY) treatment on inflammation-related gene expression. *Lateolabrax japonicus* brain (LJB) cells were transfected with 1 µg of *pCMV-Flag-CP* or *pCMV* vector for 24 h, followed by treatment with Bay 11-7082 (10 and 20 µM) for 6 h. mRNA levels of *p65*, *p100*, *p105*, *NLRP3*, *IL-1β*, and *IL-11β* were quantified by qPCR. (E) Effect of MAVS knockdown on inflammation-related gene expression. LJB cells were transfected with 1 µg of *pCMV-Flag-CP* or *pCMV* vector for 12 h, followed by transfection with either NC or MAVS siRNA for an additional 12 h. Transcript levels of *p65*, *p100*, *p105*, *NLRP3*, *IL-1β*, and *IL-11β* were measured by qPCR. All experiments were independently repeated three times. Data are shown as mean ± SD. Statistical significance was evaluated by Student’s *t*-test: *p* < 0.05 (*), *p* < 0.01 (**).

To investigate the functional consequences of CP-induced NF-κB activation, we examined the expression of NF-κB-regulated proinflammatory genes (*p65*, *p100*, *p105*, *NLRP3*, *IL-1β*, *IL-11β*). CP markedly upregulated these transcripts, while the NF-κB inhibitor Bay 11-7082 reversed these effects (Fig 1D), further confirming the involvement of NF-κB signaling in CP-induced inflammatory responses. Moreover, siRNA-mediated silencing of MAVS led to a dose-dependent reduction in CP-induced proinflammatory gene expression (Fig 1E), highlighting the critical role of MAVS in CP-mediated NF-κB activation. Collectively, these findings confirm that NNV CP induces NF-κB signaling activation and inflammatory responses, which are associated with MAVS.

### CP promotes MAVS stabilization and aggregation via SUMOylation

Given that MAVS protein stability and aggregation are critical for its signaling function [21], we next investigated the impact of CP on MAVS. Firstly, we examined whether CP affects MAVS protein stability. Co-expression of MAVS with increasing amounts of CP in HEK 293T cells resulted in a dose-dependent accumulation of MAVS protein (Fig 2A). Since SUMOylation is known to enhance protein stability, we tested the effect of the SUMOylation inhibitor Ginkgolic acid. As shown in Fig 2B, while CP overexpression elevated MAVS protein levels, Ginkgolic acid treatment resulted in a dose-dependent reduction in MAVS protein levels, suggesting that CP stabilizes MAVS via SUMOylation.

**Fig 2.**
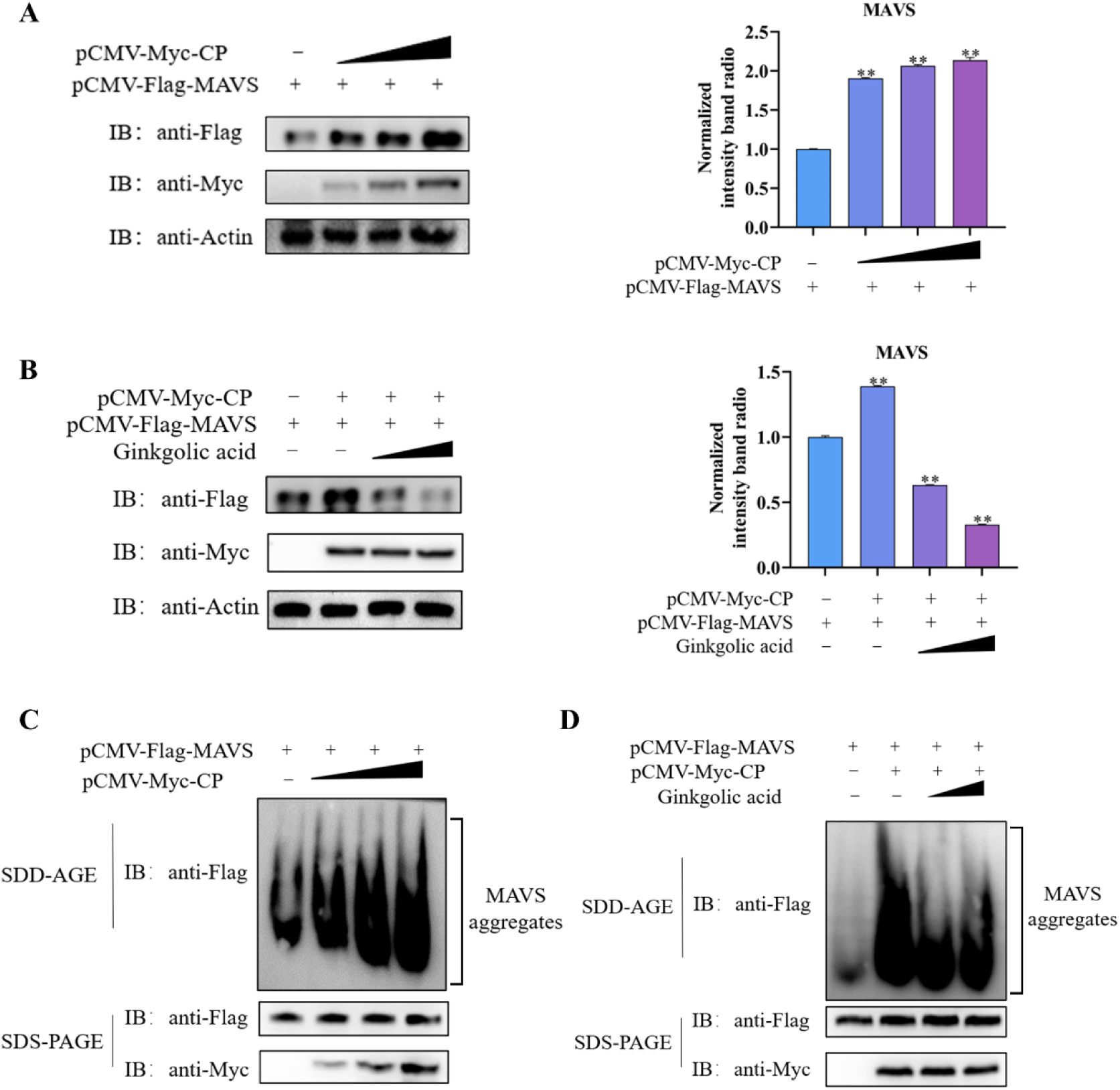
CP enhances MAVS protein levels and aggregate formation through SUMOylation. (A) Effect of CP on MAVS protein expression. HEK 293T cells were co-transfected with *pCMV-Flag-MAVS* and varying amounts of *pCMV-Myc-CP* plasmid for immunoblotting assays. Protein levels of MAVS were assessed by immunoblotting, normalized to β-actin, and quantified by densitometry (n = 3). (B) Effect of Ginkgolic acid on CP-induced MAVS protein expression. HEK 293T cells were co-transfected with *pCMV-Flag-MAVS* and *pCMV-Myc-CP* plasmids, followed by treatment with different concentrations of Ginkgolic acid (50 and 100 μM) for 6 h (DMSO as control). MAVS protein levels were analyzed by immunoblotting assays, normalized to β-actin, and quantified by densitometry (n = 3). (C) Effect of CP on MAVS aggregate formation. HEK 293T cells were co-transfected with *pCMV-Flag-MAVS* and *pCMV-Myc-CP* plasmids (0, 1, 2, or 3 µg) and treated with varying concentrations of Ginkgolic acid (50 and 100 μM) for 6 h (DMSO as control). MAVS aggregates were analyzed using semi-denaturing detergent agarose gel electrophoresis.

Next, we examined the effect of CP on MAVS aggregation, a process essential for MAVS-mediated signaling. SDD-AGE assays in HEK 293T cells co-transfected with MAVS and increasing amounts of CP revealed a dose-dependent increase in MAVS aggregates (Fig 2C). Moreover, treatment with Ginkgolic acid significantly diminished the formation of MAVS aggregates, indicating that SUMOylation is crucial for CP-induced MAVS aggregation (Fig 2D). Interestingly, confocal microscopy showed that CP did not affect the binding of MAVS to mitochondria (Fig 3), suggesting that CP-induced aggregation occurs independently of MAVS subcellular localization. These findings suggest that CP might promote MAVS aggregation via SUMOylation-mediated stabilization rather than its subcellular localization.

**Fig 3.**
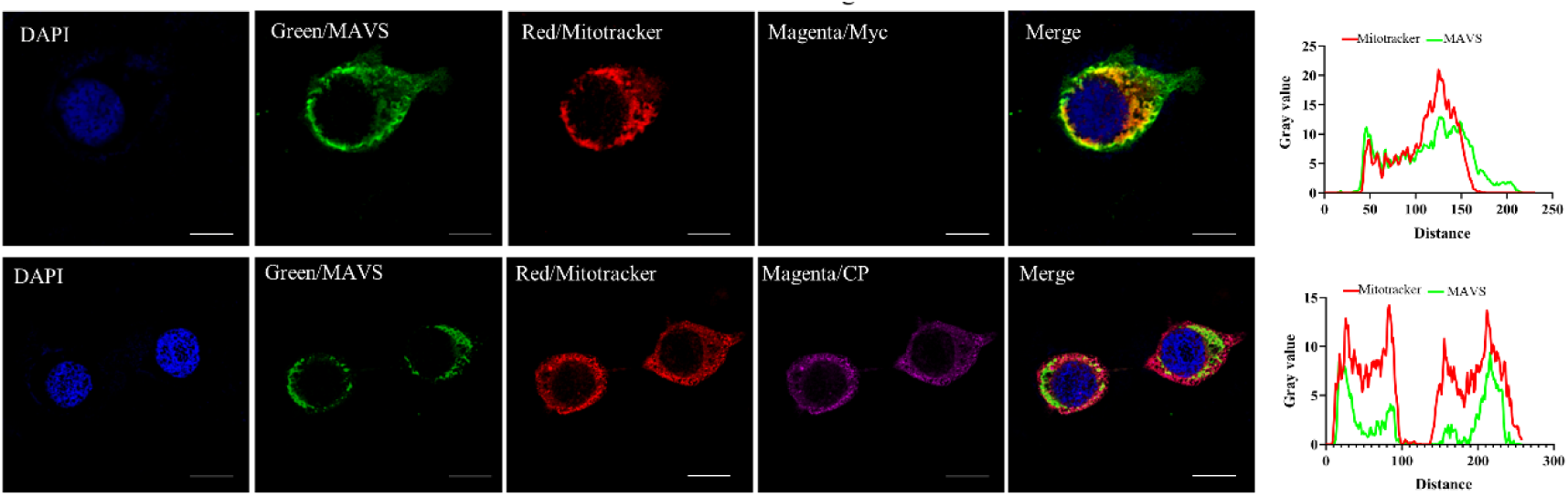
CP does not interfere with MAVS localization to mitochondria. HEK 293T cells were co-transfected with *pCMV-Flag-MAVS* and *pCMV-Myc-CP* plasmids, and analyzed by immunofluorescence (IF) assays. CP was detected using anti-Myc antibody (magenta) and MAVS with anti-Flag antibody (green). Mitochondria were labeled with MitoTracker Red (red), and nuclei were stained with DAPI (blue). Fluorescence intensity profiles were quantified using ImageJ across representative confocal fields. Scale bar = 10 µm.

### CP selectively enhances MAVS-SUMO2 interaction

To investigate the underlying mechanisms by which CP facilitates MAVS SUMOylation, we first performed a bioinformatics analysis, predicting three potential SUMOylation sites on MAVS (K47, K325, K392) within multiple SUMO-interacting motifs (Fig 4A). Confocal microscopy confirmed the cytoplasmic colocalization of MAVS with SUMO1, SUMO2, and SUMO3 (Fig 4B). Co-immunoprecipitation (Co-IP) assays demonstrated that MAVS interacted with all three SUMO isoforms, confirming its SUMOylation (Fig 4C). To assess the role of CP in MAVS SUMOylation, HEK 293T cells were co-transfected with MAVS, CP, and SUMO1/2/3 constructs. Co-IP assays revealed that CP selectively enhanced the association of MAVS with SUMO2 but not SUMO1 or SUMO3 (Fig 4D), suggesting that SUMO2 is the preferred isoform facilitating CP-mediated MAVS SUMOylation.

**Fig 4.**
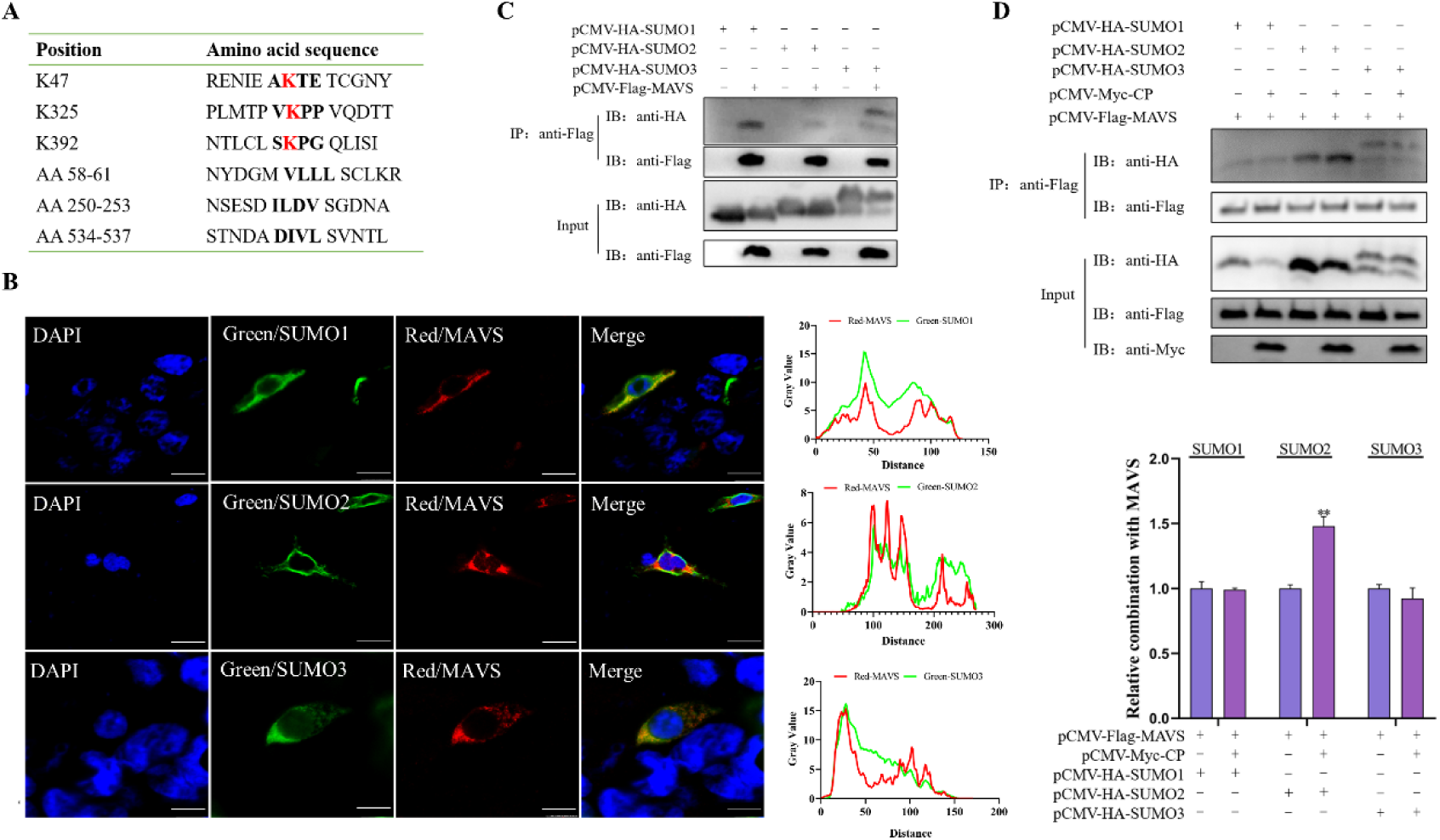
Interrelationship Between MAVS and SUMO Proteins. (A) Prediction of SUMOylation sites and SUMO-interacting motifs (SIMs) in MAVS. Potential SUMOylation sites in MAVS were predicted using the SUMOplot™ Analysis Program (Abcepta, https://www.abcepta.com/sumoplot). (B) Subcellular co-localization of MAVS with SUMO isoforms. HEK 293T cells were co-transfected with *pCMV-Flag-MAVS* and *pCMV-HA-SUMO1/2/3* plasmids, and proteins co-localization was analyzed by IF assays using anti-HA (green), anti-Flag (red), and the nucleus were stained with DAPI (blue). Co-localization signals were visualized by confocal microscopy and quantified using ImageJ. Scale bar = 10 µm. (C) Interaction between MAVS and SUMO1/2/3. HEK 293T cells were co-transfected with *pCMV-Flag-MAVS* and *pCMV-HA-SUMO1/2/3* plasmids. Co-IP assay was performed using anti-Flag magnetic beads, followed by immunoblotting to detect associated SUMO proteins. (D) CP enhances MAVS SUMOylation by SUMO2. HEK 293T cells were co-transfected with *pCMV-Flag-MAVS*, HA-tagged SUMO1/2/3, and either *pCMV-Myc-CP* or control vector. Co-IP was performed using anti-Flag magnetic beads. Bound proteins were analyzed by immunoblotting, and signal intensities were quantified by densitometry (n = 3). **, *p* < 0.01.

### CP acts as a viral SUMO E3 ligase, facilitating MAVS SUMOylation

Next, we investigated whether CP functions as a SUMO E3 ligase for MAVS. Bioinformatic analysis identified four potential SUMO-interaction motifs in CP (Fig 5A). CP overexpression alone led to time- and dose-dependent increases in *Lateolabrax japonicus* UBC9 (LjUBC9) protein levels, an E2 SUMO-conjugating enzyme (Fig 5B and 5C). Furthermore, co-transfection experiments demonstrated that CP elevated intracellular levels of SUMO1, SUMO2, and SUMO3 (Fig 5D), suggesting that CP promotes intracellular SUMOylation. Confocal microscopy revealed extensive cytoplasmic colocalization between CP and SUMO1/2/3 (Fig 6A), and Co-IP assays confirmed CP’s interaction with SUMO1, SUMO2, and SUMO3 (Fig 6B and 6C).

**Fig 5.**
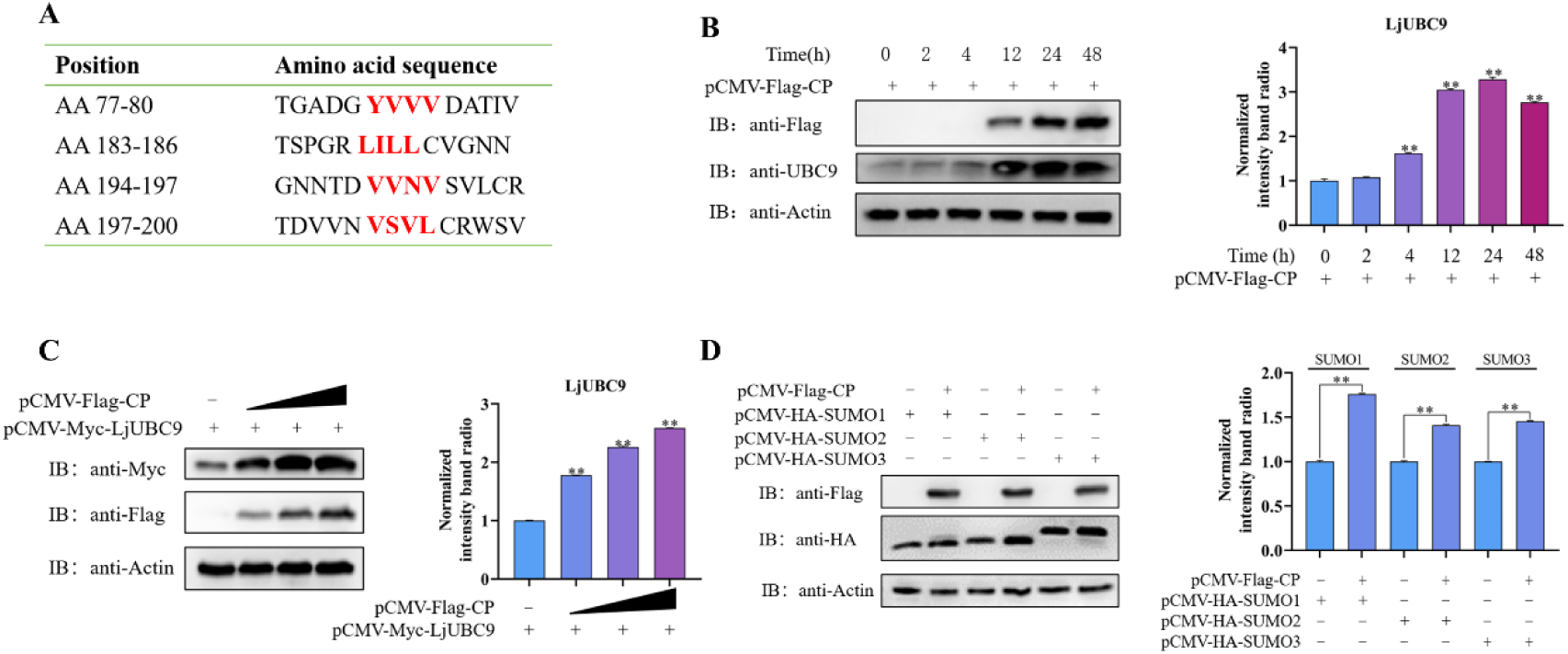
CP promotes the expression of SUMOylation pathway-related proteins. (A) Prediction of SIMs in CP. Potential SIMs in CP were predicted using the JASSA v4 web server (http://www.jassa.fr/index.php?m=jassa). (B) Time-dependent effects of CP on UBC9 protein expression. LJB cells were transfected with *pCMV-Myc-CP* and harvested at 0, 2, 4, 12, 24, and 48 h post-transfection. Protein levels of CP, LjUBC9, and β-actin were analyzed by immunoblotting. LjUBC9 expression was normalized to β-actin and quantified by densitometry (n = 3). (C) Dose-dependent effects of CP on LjUBC9 protein expression. LJB cells were transfected with increasing amounts of *pCMV-Myc-CP* (0, 1, 2, or 3 μg). Protein expression of LjUBC9 and CP was assessed by immunoblotting and quantified relative to β-actin (n = 3). (D) Effects of CP on SUMO 1/2/3 protein expression. HEK 293T cells were co-transfected with *pCMV-Flag-CP* and *pCMV-HA-SUMO1/2/3* plasmids. Protein levels were detected by immunoblotting using anti-Flag, anti-HA, and anti-actin antibodies, and quantified by densitometry (n = 3). **, *p* < 0.01.

**Fig 6.**
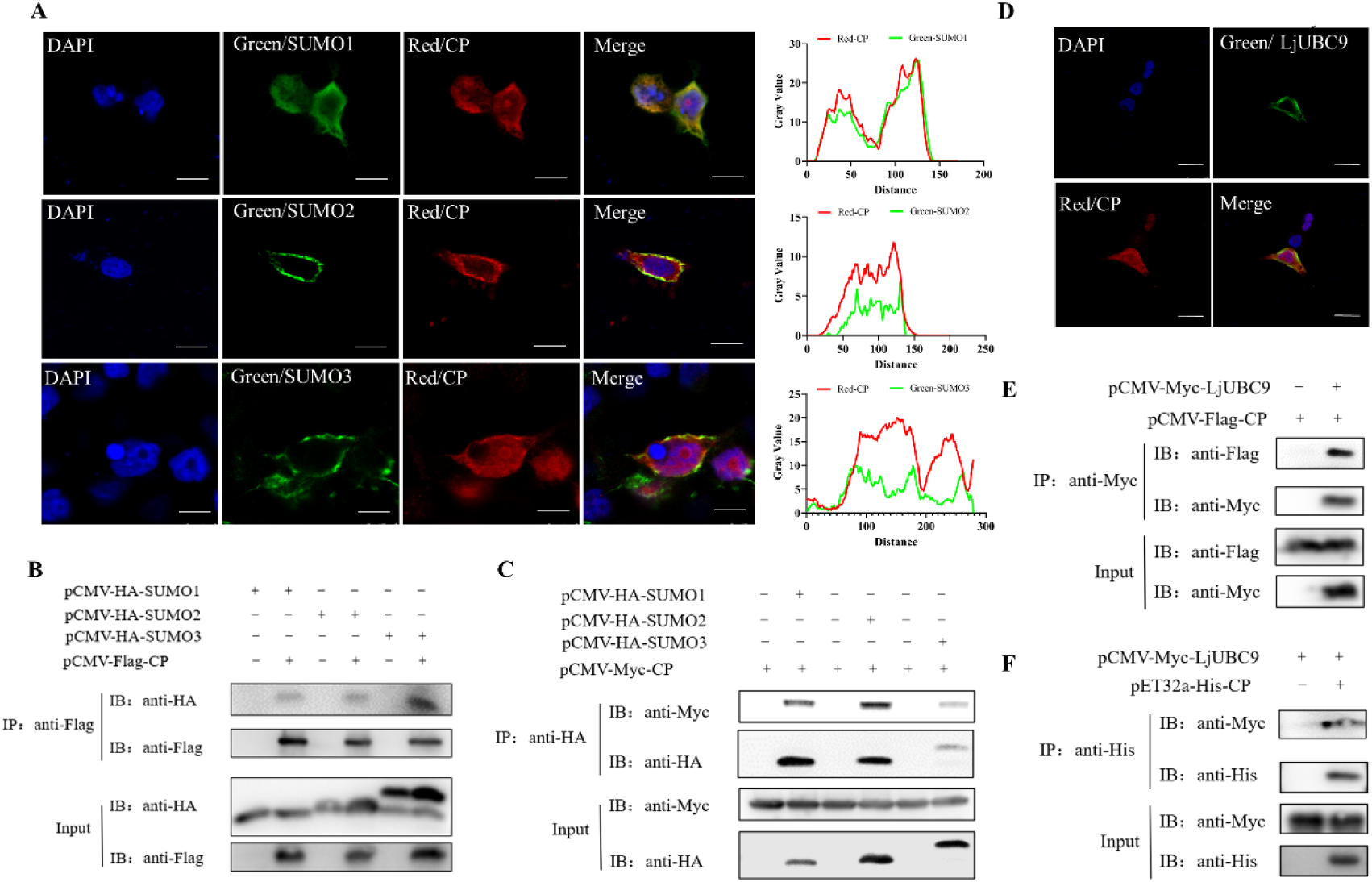
CP interacts with SUMO isoforms and the E2-conjugating enzyme LjUBC9. (A) Subcellular co-localization of CP with SUMO1, SUMO2, and SUMO3. HEK 293T cells were co-transfected with *pCMV-Flag-CP* and *pCMV-HA-SUMO1/2/3* plasmids for IF assays, using anti-HA (green), anti-Flag (red), and DAPI (blue) for nuclear staining. Co-localization signals were visualized by confocal microscopy and quantified using ImageJ. Scale bar = 10 µm. (B-C) Physical interaction between CP and SUMO isoforms. HEK 293T cells were co-transfected with *pCMV-Flag-CP* or *pCMV-Myc-CP* and *pCMV-HA-SUMO1/2/3* plasmids. Co-IP was performed using anti-Flag magnetic beads (B) or anti-HA magnetic beads (C), followed by immunoblotting. (D) Subcellular co-localization of CP with LjUBC9. HEK293T cells were co-transfected with *pCMV-Flag-CP* and *pCMV-Myc-LjUBC9* plasmids. IF assays were conducted using anti-Myc (green), anti-Flag (red), and DAPI (blue). Confocal imaging and ImageJ quantification were used to assess co-localization. Scale bar = 10 µm. (E) Co-IP analysis of the CP–LjUBC9 interaction. HEK 293T cells were co-transfected with *pCMV-Flag-CP* and *pCMV-Myc-LjUBC9* plasmids. Lysates were immunoprecipitated with anti-Flag magnetic beads and analyzed by immunoblotting. (F) Pull-down analysis of CP and LjUBC9 protein interaction. HEK 293T cells were transfected with *pCMV-Myc-LjUBC9* plasmids and pull-down assays were performed with pET32a-His-CP protein lysates. Lysates were incubated with anti-His magnetic beads for immunoprecipitation. Associated proteins were detected by immunoblotting.

Critically, CP was found to directly interact with LjUBC9, as demonstrated by confocal microscopy (Fig 6D), Co-IP and Pull-down assays (Fig 6E and 6F), fulfilling the tripartite requirement (E1-E2-E3) for SUMO ligase activity. These results suggest that CP may function as a viral SUMO E3 ligase, potentially facilitating the SUMOylation of MAVS during RGNNV infection.

### CP-mediated MAVS SUMOylation at K325 site for NF-κB activation

To identify the specific MAVS lysine residue involved in SUMOylation by CP, we generated MAVS mutants at three predicted SUMOylation sites: K47R, K325R, and K392R (Fig 7A). Co-IP assays revealed that CP preferentially SUMOylates MAVS at K325, as the MAVS-K325R mutant failed to interact with SUMO2, while MAVS-K47R and MAVS-K392R mutants retained SUMO2 binding (Fig 7B).

**Fig 7.**
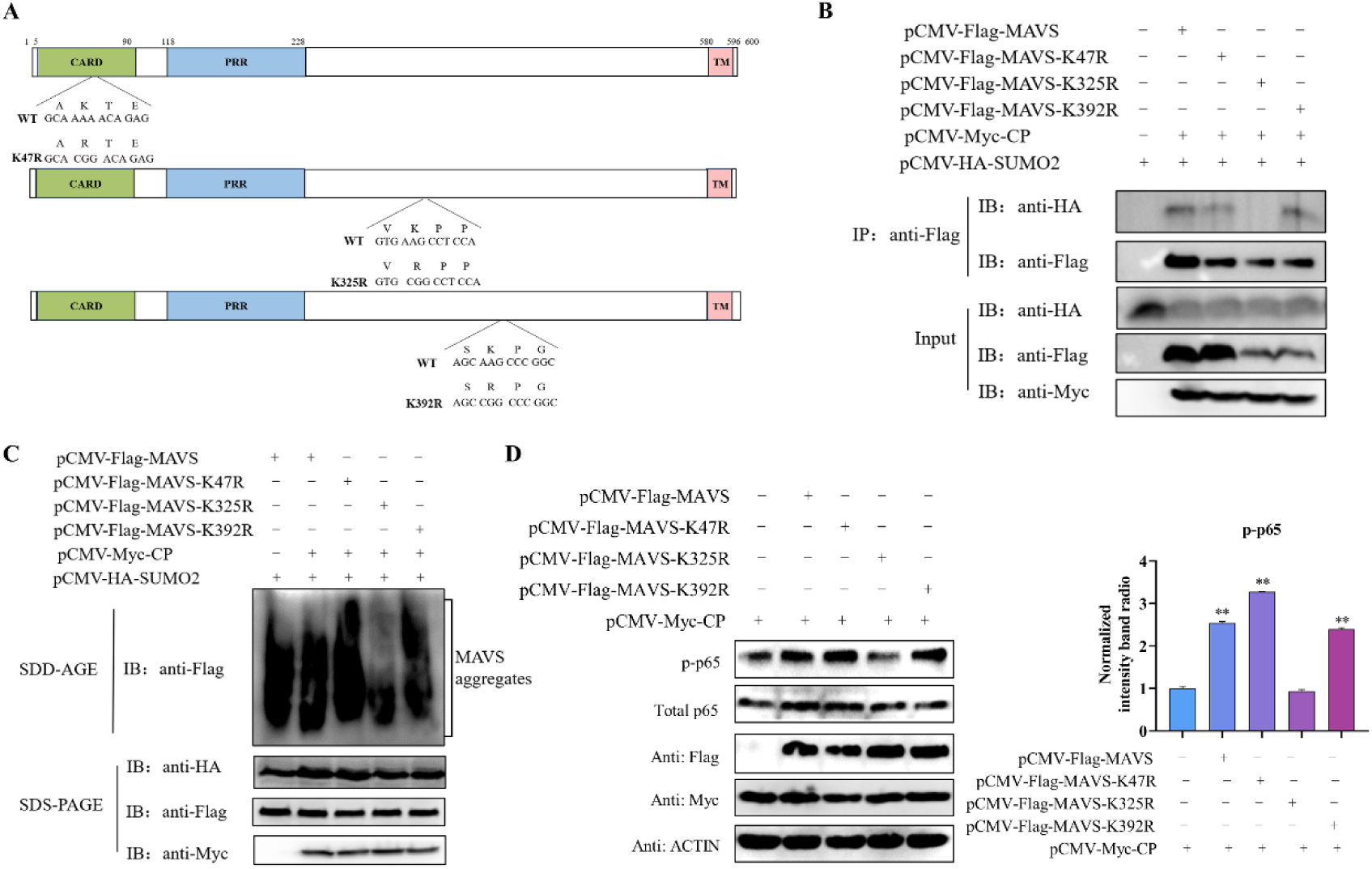
CP promotes SUMOylation of MAVS at K325. (A) Schematic representation of MAVS lysine-to-arginine mutants. The diagram illustrates the MAVS mutants with specific lysine mutations: MAVS-K47R, MAVS-K325R, and MAVS-K392R. (B) Effect of CP on the interaction between MAVS mutants and SUMO2. HEK 293T cells were co-transfected with *pCMV-HA-SUMO2*, *pCMV-Myc-CP*, and wild-type or mutant MAVS constructs (K47R, K325R and K392R). Cell lysates were immunoprecipitated using anti-Flag magnetic beads and analyzed by immunoblotting (C) Effect of K325 mutation on CP-mediated MAVS aggregation. HEK 293T cells were co-transfected with *pCMV-Myc-CP*, *pCMV-HA-SUMO2*, and *pCMV-Flag-MAVS* or MAVS site-specific mutant plasmids (K47R, K325R, and K392R) for SDD-AGE assays to assess aggregate formation. (D) Effect of CP on p65 expression induced by MAVS mutants. HEK 293T cells were co-transfected with *pCMV-Myc-CP* and wild-type or mutant MAVS plasmids. Protein levels of p65 and p-p65 were assessed by western blotting, normalized to β-actin, and quantified by densitometry (n = 3). **, *p* < 0.01.

To examine the functional consequences of the K325 mutation, we analyzed MAVS aggregation and NF-κB activation. The MAVS-K325R mutant failed to form aggregates in response to CP overexpression (Fig 7C). Furthermore, CP-mediated activation of NF-κB was abolished in cells expressing MAVS-K325R, as evidenced by the lack of p65 accumulation (Fig 7D). These results confirm that K325 is the key residue for CP-induced MAVS SUMOylation, aggregation, and subsequent NF-κB activation.

## Discussion

The innate immune system plays a pivotal role in defending against viral infections, with MAVS serving as a central mediator in antiviral signaling. MAVS transduces signals from RLRs to activate downstream antiviral pathways, including NF-κB and IFN signaling. While much has been understood about MAVS’s role in immune responses, the mechanisms by which viruses manipulate MAVS function remain incompletely explored. NNV is a potent pathogen that causes severe viral infections in fish, triggering robust immune responses in the host. While previous studies have shown that NNV infection activates NF-κB signaling [15], the molecular mechanisms underlying this activation remained unclear. In this study, we reveal a novel mechanism by which NNV, through its CP, modulates the MAVS-NF-κB signaling axis. Our findings demonstrate that NNV CP acts as a viral SUMO E3 ligase, facilitating the SUMOylation of MAVS at K325, which enhances MAVS stabilization and aggregation and activates the NF-κB signaling pathway (Fig 8). This represents a previously unrecognized strategy by which NNV exploits the host SUMOylation machinery to modulate immune responses and facilitate viral persistence.

**Fig 8.**
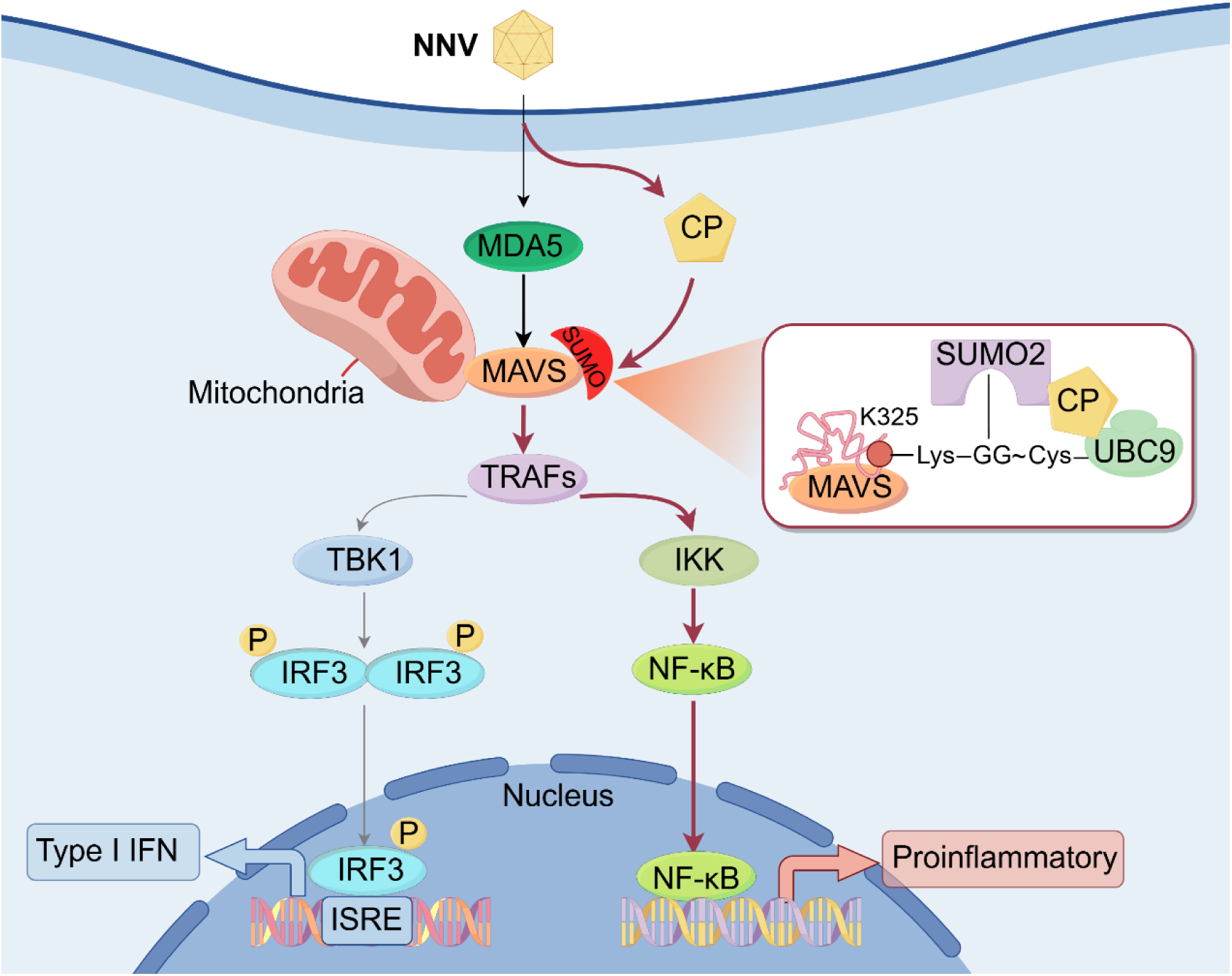
Schematic description of CP exploits SUMOylation to amplify MAVS-mediated NF-κB signaling. CP strengthens the binding between MAVS and SUMO2 and enhances SUMOylation of MAVS via acting as a viral SUMO E3 ligase. The SUMO-modified MAVS exhibits increased aggregation, which in turn amplifies NF-κB signaling.

SUMOylation is a vital post-translational modifications (PTM) in eukaryotes, regulating various cellular processes including signal transduction, protein localization, and transcriptional activity [22]. Viruses have evolutionarily exploited this system to enhance replication and immune evasion, with many hijacking host SUMOylation machinery to modify key immune components [23]. Our study reveals that NNV CP functions as a viral-encoded SUMO E3 ligase to catalyze MAVS SUMOylation. Experimental evidence demonstrates three synergistic mechanisms underlying CP’s activity: First, bioinformatics analysis predicts potential SUMO interaction motifs within the CP sequence, suggesting a potential role for CP in SUMOylation. Second, direct interaction between CP and the SUMO E2 enzyme LjUBC9 [24] is validated by Co-IP and pull-down assays, fulfilling the E2-E3 complex requirement for SUMOylation. Additionally, confocal microscopy and Co-IP assays demonstrate that CP interacts with SUMO isoforms, suggesting spatially coordinated modification. These findings are further supported by SUMOylation inhibitor studies: Ginkgolic acid [25] dose-dependently reverses CP-induced MAVS stabilization, confirming enzymatic dependency. The ability of viral proteins to hijack the host’s SUMOylation machinery for immune modulation is not unique to NNV. Similarly, the presence of viral proteins functioning as SUMO ligases has also been observed in other viral species, which target host proteins via SUMOylation to modulate immune responses. For example, the ORF45 of kaposi’s sarcoma-associated herpesvirus acts as a SUMO E3 ligase, regulating the 90-kDa ribosomal S6 kinases 1 SUMOylation to promote viral lytic replication [26]. The adenovirus E4-ORF3 acts as a SUMO E3 ligase, promoting transcriptional intermediary factor 1 gamma SUMOylation and poly-SUMO chain elongation to regulate cellular processes during infection [27].

MAVS is a critical adaptor protein that integrates signals from RLRs to initiate antiviral responses. As such, its activation and stability are tightly regulated by multiple PTMs, including ubiquitination, phosphorylation, and SUMOylation [28]. These PTMs ensure that MAVS operates at the appropriate intensity and duration during viral infections. For example, the E3 ubiquitin ligase TRIM25 facilitates the K7- and K10-linked polyubiquitination of MAVS, which is crucial for its activation [29]; whereas RNF125 mediates K48-linked polyubiquitination, targeting MAVS for proteasomal degradation, thus controlling the threshold of MAVS activation and limiting excessive immune responses [30]. In human keratinocytes, MAVS is found to interact with SUMO3 and undergo SUMOylation, which is essential for the activation of IFN-β secretion during antiviral responses [9]. During Sendai virus infection, SENP1 removes poly-SUMO chains from MAVS and inhibits MAVS aggregation [10]. The RIG-I ligand poly(dA:dT) was also found to promote the binding of MAVS to SUMO3, thereby enhancing MAVS aggregate formation [9]. Our previous study demonstrated that *L. japonicus* RNF114 mediates K27- and K48-linked polyubiquitination of MAVS, leading to its degradation and suppression of the IFN response [18], among which, CP was found inducing the expression of RNF114, but antagonizes RNF114-induced MAVS degradation, suggesting that CP might manipulate MAVS expression through a mechanism other than ubiquitination. Here, we demonstrate that CP promotes MAVS stability and aggregation through SUMOylation. This SUMOylation-mediated stabilization of MAVS represents a novel viral strategy for modulating host immune signaling. Moreover, we found the selective enhancement of MAVS-SUMO2 interaction by CP. This preference for SUMO2 over other isoforms (SUMO1 and SUMO3) suggests that poly-SUMO chains, which are more commonly associated with SUMO2, play a crucial role in MAVS aggregation and signaling [31]. In contrast, SUMO1 is typically involved in mono-SUMOylation, which may have distinct regulatory functions [24]. This selective modification is likely a crucial determinant in the ability of CP to amplify MAVS function, as poly-SUMO chains are known to recruit effector proteins that enhance signal transduction.

MAVS modification at specific lysine residues regulates its stability, localization, and activation of downstream pathways. The balance between ubiquitination-mediated degradation and SUMOylation-induced stabilization is influenced by these modification sites, underscoring a complex regulatory interplay [23]. Research has suggested that certain lysine residues, such as K10, K311, K461 of MAVS, may be important for TRIM31 mediated K63-linked ubiquitylation [32]. K48-linked ubiquitylation targets MAVS at K7, K10, K371, K420 and K500 sites for proteasomal degradation limits the activation of MAVS [33]. K27-linked ubiquitylation induced by TRIM21 targets at K325 to induce the recruitment of TBK1 to MAVS, facilitating the activation of downstream antiviral signaling pathways [5]. Although the study of MAVS SUMOylation sites is still not sufficiently thorough, it suggests SUMOylation modification of MAVS typically occurs at lysine residues within the consensus motif ΨKxD/E [7]. Of note, MAVS K461 and K500 not only act as the ubiquitylation sites [32, 33], but also serve as SUMOylation sites mediated by PIAS3, which induce MAVS aggregation and phase separation, suggests a dual role of K461 and K500 as sites for both SUMOylation and ubiquitylation in fine-tuning MAVS activity [10]. We found that the K325 residue on MAVS is critical for its SUMOylation, aggregation, and subsequent activation of NF-κB signaling, which highlights the complex regulation of MAVS and its multifaceted role in innate immunity. Given that *L. japonicus* RNF114 mediates K27- and K48-linked polyubiquitination of MAVS, future research should focus on investigating how SUMOylation interacts with other PTMs, such as ubiquitination and phosphorylation, in regulating MAVS function.

The activation of NF-κB during viral infections is widely recognized as advantageous for viruses, as it can prevent apoptosis, promote inflammation, and modulate immune responses to favor viral survival and replication [34, 35]. MAVS aggregation is sufficient to trigger NF-κB signaling, which can lead to either the production of type I IFNs (a key antiviral response) or the release of pro-inflammatory cytokines that may exacerbate immunopathology [3, 36]. This functional duality reveals a delicate “immunological balance” wherein viruses benefit from moderate NF-κB activation that sustains inflammation conducive to replication, while concurrently suppressing IFN responses to evade antiviral defenses [37, 38]. For example, SARS-CoV-2 ORF3a activates NF-κB to promote IL-6 production, creating a pro-inflammatory microenvironment that enhances viral spread while potentially exacerbating immunopathology [39]. HCV cleaves MAVS via NS3/4A protease, inhibiting IFN while sustaining controlled NF-κB activation to prevent apoptosis [40, 41]. In the case of NNV, the CP appears to employ a comparably nuanced strategy to manipulate host immunity. Previous studies have shown that NNV CP suppresses IFN signaling by promoting MAVS degradation [18]. Here, our findings further reveal that NNV CP modulates MAVS through opposing PTMs: it facilitates MAVS ubiquitination to suppress IFN responses and simultaneously induces MAVS SUMOylation to sustain NF-κB activation. This dual mechanism enables NNV to suppress the host’s antiviral defense while leveraging the inflammatory response for its own replication advantage. The coexistence of these two PTMs:ubiquitination leading to IFN suppression and SUMOylation promoting NF-κB-driven inflammation, underscores NNV’s evolved capacity to fine-tune host immune pathways. By differentially regulating MAVS, NNV CP achieves a balance that favors viral survival: evading immune clearance while exploiting inflammation to enhance replication and spread.

In conclusion, our study identifies NNV CP as a key viral modulator of the host immune response, functioning as a SUMO E3 ligase to promote SUMOylation of MAVS. This modification enhances MAVS aggregation, stabilizes its function, and activates the NF-κB signaling pathway, contributing to the inflammatory response during viral infection. This work provides important new insights into the PTM of MAVS, highlighting the potential of targeting SUMOylation pathways as a strategy to combat viral infections and associated immune dysregulation.

## Materials and methods

### Cell, virus, and reagents

*Lateolabrax japonicus* brain cell line (LJB) was originally established in our laboratory and cultivated in Dulbecco’s modified Eagle medium (DMEM) incorporating 15% (v/v) fetal bovine serum (FBS) and 1% (g/v) basic fibroblast growth factor at 28℃, as previously described [42]. Fathead minnow (FHM) cells were cultivated in Medium 199 containing 10% FBS at 28 ℃. Human embryonic kidney 293T (HEK 293T) cells were sustained in DMEM supplemented with 10% FBS at 37 °C in a humidified atmosphere with 5% CO₂.

Red-spotted grouper nervous necrosis virus (RGNNV) was isolated from infected *L. japonicus* collected from an aquaculture farm in Guangdong Province, China. The virus was propagated in LJB cells and stored at −80 °C until use [43].

Primary antibodies included rabbit polyclonal anti-UBC9 (T55571), mouse monoclonal anti-Myc (M20002), anti-HA (M20003), anti-Flag (M20008), anti-β-actin (M20011), and anti-His (M30111), all obtained from Abmart (Shanghai, China). Rabbit polyclonal anti-Flag (YM3201) and anti-Myc (YM3203) antibodies were purchased from ImmunoWay (Texas, USA).

The following reagents were obtained from MedChemExpress (Shanghai, China): ginkgolic acid (HY-N0077), NF-κB inhibitor Bay 11-7082 (HY-13453), protease inhibitor cocktail (HY-K0010), and magnetic beads conjugated with anti-Flag (HY-K0207), anti-c-Myc (HY-K0206), anti-HA (HY-K0201), or anti-His (HY-K0209) antibodies. Horseradish peroxidase (HRP)-conjugated goat anti-mouse IgG (H+L) (A0216) and goat anti-rabbit IgG (H+L) (A0208), cell lysis buffer (P0013), Lipofectamine™ 8000 (C0533), and His-tag protein purification kit (P2226) were obtained from Beyotime (Shanghai, China).

Fluorescent secondary antibodies, including Alexa Fluor 488-conjugated donkey anti-mouse IgG (A21202) and Alexa Fluor 555-conjugated donkey anti-rabbit IgG (A31572), were purchased from Sigma-Aldrich (Darmstadt, Germany). DAPI staining solution (40728ES03) was obtained from Yeasen Biotechnology (Shanghai, China). TRIzol reagent, pCMV-Myc, pCMV-Flag, and pCMV-HA vectors were acquired from Invitrogen (California, USA). Immobilon Western Chemiluminescent HRP Substrate (WBKLS0500) was obtained from Merck Millipore (Darmstadt, Germany). One-step PAGE gel fast preparation kit (E303-01) and 2× Phanta Flash Master Mix (P520) were purchased from Vazyme (Nanjing, China). Restriction enzymes *EcoRI-HF* (R3101), *KpnI-HF* (R3142), *SalI* (R0138), and *XhoI* (R0146) were sourced from New England Biolabs (Beijing, China).

### Plasmids construction

The coding sequence of SUMO1, SUMO2, SUMO3 were cloned into the *KpnⅠ* and *SalⅠ or EcoRⅠ* sites of *pCMV-HA* vector according to molecular biology techniques. Site-directed mutagenesis was performed on the pCMV-Flag-MAVS plasmid to generate lysine-to-arginine mutants (K47R, K325R, and K392R), resulting in pCMV-Flag-MAVS-K47R, -K325R, and -K392R constructs. All sequences were verified by DNA sequencing. Primer sequences are listed in S1 Table.

The plasmids pCMV-Myc-CP, pET32a-His-CP, pCMV-Flag-MAVS, and pGL3-NF-κB-Luc-pro were preserved in our laboratory.

### Cell Transfection and RNA Interference (RNAi)

For MAVS gene silencing, specific siRNAs were designed and commercially synthesized by Tsingke Biotechnology Company (siRNA sequences were provided in S2 Table). Transfection efficiency was assessed by qRT-PCR, and the optimal siRNA (MAVS-siRNA2: 5′-GCUUCUUCUGAGCUGUCUA(dT)(dT)-3′) showing the highest knockdown efficacy was chosen for further studies.

Prior to transfection, cells were seeded in culture plates and allowed to adhere overnight until reaching 80%–90% confluence. Transfection was carried out using Lipofectamine™ 8000 transfection reagent (Beyotime, Shanghai, China) according to the manufacturer’s protocol and previous reports [18]. Briefly, cells in 6-well plates were transfected with 100 nM MAVS siRNA or negative control (NC) siRNA using 4 μL of transfection reagent. After 4–6 h of incubation, the medium was replaced with fresh culture medium. Cells were harvested at 24 h post-transfection for qRT-PCR analyses.

### Dual luciferase reporter activity assays

Dual-luciferase reporter assays were performed as previously described using the Dual-Luciferase Reporter Assay System (Promega) [18]. FHM cells were seeded in 24-well plates and transfected with the firefly luciferase reporter plasmid (pGL3-NF-κB-pro-Luc), internal control plasmid pRL-TK (Promega), and other indicated plasmids at a ratio of 50:1:50. After 24 h, cells were lysed with Passive Lysis Buffer and luminescence was measured using a GloMax 20/20 luminometer (Promega). Firefly luciferase activity was normalized to Renilla luciferase activity to account for transfection efficiency. Data are presented as mean ± standard deviation (SD) from three independent experiments.

### Inhibitors treatment

Ginkgolic acid and BAY 11-7082 were dissolved in dimethyl sulfoxide (DMSO) to a stock concentration of 10 mM and stored at −80 °C. For treatment, Ginkgolic acid was diluted in culture medium to final concentrations of 50 μM and 100 μM, while BAY 11-7082 was used at 10 μM and 20 μM. Cells were treated with these inhibitors for 6 h, with DMSO used as a vehicle control.

### Confocal laser microscopy assays

Confocal microscopy was performed as previously described [44]. HEK 293T cells were seeded on cell climbing coverslips in 24-well plates and co-transfected with the indicated plasmids. At 24 h post-transfection, cells were washed three times with PBS, fixed with 4% paraformaldehyde for 10 min at room temperature, and permeabilized with 0.1% Triton X-100 in PBS. After blocking with 5% skim milk in PBS for 1 h, cells were incubated overnight at 4 °C with anti-Flag primary antibody (1:400). After washing with PBS, cells were incubated for 1 h at room temperature with Alexa Fluor 555-conjugated goat anti-rabbit IgG (1:400) and DAPI (1:1000) for nuclear staining. Images were captured using a ZEISS laser scanning confocal microscope.

### Bioinformatics analysis

Potential SUMOylation sites in MAVS and SUMO-interacting motifs (SIMs) in CP were predicted using the SUMOplot™ Analysis Program (Abcepta, https://www.abcepta.com/sumoplot) and the JASSA v4 web server (http://www.jassa.fr/index.php?m=jassa).

### Co-immunoprecipitation (Co-IP) and immunoblot (IB) assays

Co-IP assays were conducted with minor modifications from our previous protocol [45]. HEK 293T cells were seeded in 60 mm dishes and co-transfected with the indicated plasmids for 48 h. Cells were washed twice with 4 mL of cold PBS, lysed in 500 μL of lysis buffer supplemented with protease inhibitor cocktail and 20 mM N-ethylmaleimide, and disrupted by sonication (3 cycles). Lysates were centrifuged at 12,000×*g* for 15 min at 4 °C, and supernatants were incubated with antibody-conjugated magnetic beads overnight at 4 °C with gentle rotation. Beads were washed five times with RIPA buffer and subjected to immunoblot analysis.

For immunoblotting, cell lysates or immunoprecipitates were separated by 10% or 15% SDS-PAGE and transferred onto 0.22 μm PVDF membranes. Membranes were blocked with 5% skim milk in TBST (25 mM Tris-HCl, 150 mM NaCl, 0.1% Tween-20, pH 7.5) for 1 h at room temperature and incubated overnight at 4 °C with primary antibodies. After washing, membranes were incubated with corresponding secondary antibodies for 1 h at room temperature. Protein bands were visualized using enhanced chemiluminescence (ECL) on a chemiluminescence imaging system.

### His-Tag fusion protein purification and pull-down assays

His-tagged CP proteins were purified from *Escherichia coli* BL21 transformed with the indicated prokaryotic expression plasmids. The bacteria were cultured in 30 mL LB medium supplemented with 0.5 mM isopropyl-β-D-thiogalactopyranoside at 18 °C, 120 rpm for 16 h. Cells were harvested by centrifugation and lysed, and His-tagged proteins were purified using His-tag Purification Resin (Beyotime) according to the manufacturer’s instructions. The purified protein lysate was used for pull-down assays.

Pull down assays were conducted as we formerly described [44]. Purified His-tagged proteins were first incubated with anti-His magnetic beads at room temperature with gentle rotation. The beads were washed three times with RIPA buffer (50 mM Tris-HCl [pH 7.4], 150 mM NaCl, 1% NP-40, 0.1% SDS, and 2 mM sodium deoxycholate), followed by incubation with lysates from HEK 293T cells transfected with the indicated plasmids at 4 °C overnight. The beads were then washed five times with RIPA buffer and subjected to immunoblot (IB) analysis.

### Semi-denaturing detergent agarose gel electrophoresis (SDD-AGE) assays

SDD-AGE was performed based on a published protocol with slight modifications [46]. HEK 293T cells transfected with the indicated plasmids were lysed in 1× cell lysis buffer (0.5× TBE, 10% glycerol, 2% SDS, 0.0025% bromophenol blue, and 20 mM N-ethylmaleimide). Protein samples were resolved on a vertical 1.5% agarose gel prepared in 1× TBE. Electrophoresis was performed at constant voltage in 1× running buffer (1× TBE containing 0.1% SDS) for 80 min in an ice bath. Proteins were then transferred to 0.22 μm PVDF membranes and analyzed by immunoblotting.

### Statistical analysis

Statistical analysis was performed using Student’s *t*-test for comparisons between two groups and one-way ANOVA for comparisons among multiple groups. Data were analyzed using SPSS 23.0, and graphs were generated using GraphPad Prism 8.0. *P* values less than 0.05 were considered statistically significant, and values less than 0.01 were considered highly significant.

## Data Availability Statement

The authors confirm that the data supporting the findings of this study are available within the article.

## Author contributions

**Conceptualization**: Meisheng Yi, Kuntong Jia.

**Data curation**: Wanwan Zhang, Xiaoqi Chen.

**Formal analysis**: Wanwan Zhang, Lan Yao.

**Funding acquisition**: Meisheng Yi, Kuntong Jia, Wanwan Zhang.

**Investigation**: Wanwan Zhang, Xiaoqi Chen, Bingbing Sun, Xingchen Xiong.

**Methodology**: Wanwan Zhang, Xiaoqi Chen.

**Project administration**: Meisheng Yi, Kuntong Jia.

**Supervision**: Meisheng Yi, Kuntong Jia.

**Visualization**: Wanwan Zhang, Xiaoqi Chen.

**Writing – original draft**: Wanwan Zhang, Xiaoqi Chen, Kuntong Jia.

**Writing – review & editing**: Wanwan Zhang, Meisheng Yi, Kuntong Jia

